# Cardiomyocyte-restricted expression of IL11 causes cardiac fibrosis, inflammation, and dysfunction

**DOI:** 10.1101/2023.05.23.541928

**Authors:** Mark Sweeney, Katie O’Fee, Chelsie Villanueva-Hayes, Ekhlas Rahman, Michael Lee, Konstantinos Vanezis, Ivan Andrew, Wei-Wen Lim, Anissa Widjaja, Paul JR Barton, Stuart A Cook

**Affiliations:** MRC-London Institute of Medical Sciences, Hammersmith Hospital Campus, London, UK; Institute of Clinical Sciences, Faculty of Medicine, Imperial College, London, UK; Wellcome Trust / NIHR 4i Clinical Research Fellow, Imperial College, London, UK; National Heart and Lung Institute, Imperial College, London, UK; National Heart Research Institute Singapore, National Heart Centre Singapore, Singapore; Cardiovascular and Metabolic Disorders Program, Duke-National University of Singapore Medical School, Singapore; Royal Brompton and Harefield Hospitals, Guy’s and St. Thomas’ NHS Foundation Trust, London UK

**Keywords:** Interleukin-11 Fibrosis EndMT, Heart failure

## Abstract

Background: Cardiac fibrosis is a common pathological process in heart disease and represents a therapeutic target. TGFβ is the canonical driver of cardiac fibrosis and was recently shown to be dependent on IL11 for its profibrotic effects in fibroblasts. In the opposite direction, recombinant human IL11 has been reported as anti-fibrotic and also anti-inflammatory in the mouse heart. Objectives: In this study, we determined the effects of IL11 expression in cardiomyocytes on cardiac pathobiology and function. Methods: We used the Cre-loxP system to generate a tamoxifen-inducible mouse with cardiomyocyte-restricted murine *Il11* expression. Using protein assays, bulk RNA-sequencing, and *in vivo* imaging we analysed the effects of IL11 on myocardial fibrosis, inflammation and cardiac function and challenge previous reports suggesting cardioprotective potential of IL11. Results: TGFβ stimulation of cardiomyocytes caused *Il11* upregulation. As compared to wild-type controls, *Il11* expressing hearts demonstrated severe cardiac fibrosis and inflammation that was associated with the upregulation of cytokines, chemokines, complement factors and increased inflammatory cells. IL11 expression also activated a programme of endothelial-to-mesenchymal transition and resulted in left ventricular dysfunction. Conclusion: Our data define species matched IL11 as strongly profibrotic and proinflammatory when secreted from cardiomyocytes and further establish IL11 as a disease factor.

## Introduction

Interleukin-11 (IL11) is a little-studied and poorly understood member of the IL-6 family of cytokines. Initial reports on the role of IL11 in the myocardium described cardioprotective, anti-fibrotic and anti-inflammatory effects of recombinant human IL11 (rhIL11) in the setting of myocardial ischaemia in the mouse.^1^ In keeping with this, studies in other organ systems have shown anti-inflammatory effects of rhIL11 in other mouse models of disease.^2^ However, a major limitation of these early studies was the systematic use of non-species-matched rhIL11 in mice and it was subsequently shown that rhIL11 has unexpected effects in the mouse.^3^ More recently, a range of loss- and (species-matched) gain-of-function studies have shown a profibrotic and proinflammatory effect of IL11 in various contexts.^4–8^

IL11 is not easily detected in healthy tissues but is expressed in parenchymal, mesenchymal and epithelial cells as an alarmin-like response following exposure to a range of cell stressors such as viral, toxin, bacterial, inflammatory, mechanical, oxidative and genetic factors.^9^ In particular, TGFβ strongly upregulates IL11 in human and mouse cardiac fibroblasts.^5^ IL11 expression is increased In mouse models of cardiac disease including transverse aortic constriction^10^, angiotensin II infusion^11^ and ischaemia^1^. In human studies, administration of exogenous IL11 causes heart failure symptoms and elevation of serum B-natriuretic peptide levels and IL11 levels are associated with atrial fibrosis.^12, 13^

To date, the effects of IL11 in the heart have focused on its role in fibroblasts using organismal-level loss- and gain-of-function studies^5^ and the effect of IL11 secretion from cardiomyocytes is not known. Equally, the true activity of IL11 in cardiac inflammation remains poorly characterised and controversial.^1, 5, 14^ In this study, we established a new transgenic mouse model with temporally regulated and cardiomyocyte-specific *Il11* expression to understand the effects of IL11 secretion from non-mesenchymal cells in the adult heart.

## Methods

### Animal studies

All mouse studies were conducted according to the Animals (Scientific Procedures) Act 1986 Amendment Regulations 2012 and approved by the Animal Welfare Ethical Review Body at Imperial College London. Mice were housed in the same room with a 12-hour light-dark cycle and provided food and water ad libitum. Female Rosa26 *Il11* mice on a C57BL/6N background were used as previously described (Jax cat: 031928)^5^ and male α-MHC-MerCreMer (MCM) on a C57BL6/J background were purchased from Jax (cat:005657).

Expression of *Il11* was induced in 8-week-old mice heterozygous for both MCM and Rosa26-*Il11* transgenes (designated as MCM^Cre/+^:R26R^Il^^11/+^; CM-Il11-Tg). Age-matched littermates, heterozygous for the MCM gene and wild-type for the Rosa26-*Il11* transgene (MCM^Cre/+^:R26R^Il^^11/+^; Controls), were used as control animals to account for Cre-mediated toxicity phenotypes previously reported with MCM mice.^15^

Tamoxifen (Sigma-Aldrich, T5648) was dissolved in ethanol (Fisher Scientific, 10437341, 10% v/v) and corn oil (Sigma-Aldrich, C8267) and administered intraperitoneally for 5 consecutive days at 20 mg/kg from 8 weeks of age and followed up at the designated time points (1-, 3- and 6-weeks post transgene induction).

### Genotyping

Mice were bred in a dedicated breeding facility with ad libitum access to food and water with a controlled 12-hour light-dark cycle. Genotype was confirmed with ear-notch DNA samples. DNA was extracted using a sodium hydroxide digestion buffer, then diluted in 1M Tris-HCl pH 8. Rosa26-*Il11* transgene genotype was confirmed with a single PCR reaction yielding a PCR product at 270bp in the wild-type or 727bp in the transgenic mouse. MCM mice were genotyped using two reactions for either the transgenic gene at 295pb or wild-type gene at 300bp along with an internal positive control. Primers used in these reactions are detailed in Supplementary Table 1.

### Cardiomyocyte extraction

Male wild-type C57BL/6J mice at 12 weeks of age were deeply anaesthetized with ketamine and xylazine before the heart was harvested. The chest was opened and the heart and aorta were transferred to ice-cold perfusion buffer (113mM NaCl, 4.7mM KCl, 0.6mM KH2PO4, 0.6mM Na2HPO4, 1.2mM MgSO4-7H2O, 12mM NaHCO3, 10mM KHCO3, HEPES Na Salt 0.922mM, Taurine 30mM, BDM 10mM, glucose 5.5mM, pH 7.4). Excess material was dissected away from the heart and the ascending aorta cannulated and placed on a retrograde perfusion Langendorff apparatus. The heart was perfused with warmed perfusion buffer for 4 mins and then digestion buffer (0.2mg/mL Liberase TM, 5.5mM Trypsin, 0.005U/mL DNase, 12.5uM CaCl2) until fully digested. Subsequently, the heart was minced and gently pipetted until a uniform suspension was formed. Cells were filtered through sterile gauze and resuspended in perfusion buffer. Sequential calcium reintroduction was performed in 3 steps up to 1mM (62uM, 212uM and 1mM). Cardiomyocytes were plated on laminin and allowed to equilibrate for 2 hours in M199 media supplemented with 2mM L-carnitine, 5mM creatine, and 5mM taurine. Blebbistatin (Sigma Aldrich, 203390) was added at 25uM to prevent cardiomyocyte contraction and extend culture life. Cells were treated with media containing recombinant mouse transforming growth factor-beta 1 (rmTGFβ1) at 5ng/mL (R&D systems, 7666-MB) or normal media. The media was removed after 24 hours, washed and RNA was harvested using TRIzol reagent (Invitrogen, 15596026).

### Echocardiography

Echocardiographic images were measured under anaesthesia using a Vevo3100 imaging system (Fujifilm Visualsonics Inc., USA). Anaesthesia was induced with 4% isoflurane and maintained with isoflurane 1.5-2%. Measurements were averaged across three heartbeats and strain analysis was performed in a semi-automated manner using VevoStrain imaging (Fujifilm Visualsonics Inc., USA) software.

### qPCR

The heart was dissected from the chest and washed in ice-cold PBS and snap-frozen in liquid nitrogen. Total RNA was extracted using TRIzol (Invitrogen, 15596026) in RNeasy columns (Qiagen, 74106). cDNA was synthesised using Superscript Vilo Mastermix (Invitrogen, 11755050). Gene expression analysis for *Il11* was performed using a Taqman probe (Mm00434162_m1) over 40 cycles. Expression of genes involved in fibrosis and extracellular matrix turnover were analysed in duplicate using a custom TaqMan array microfluidic card. 100µL of sample cDNA and master mix was added to each well and the card was centrifuged twice for 1 minute at 331 x g and analysed in duplicate over 40 cycles. Primers and probes used are detailed in Supplementary Table 2. Expression data were normalised to *Gapdh* mRNA expression and fold change compared to control samples was calculated using 2-ΔΔCt method.

### RNASeq

RNA quality for RNAseq was assessed using the Agilent 2100 RNA 6000 Nano assay and RNASeq libraries prepared using the NEBNext Ultra II Directional RNA Library Prep Kit with NEBNext Poly(A) mRNA Magnetic Isolation Module following manufacturer’s instructions.

Library quality was evaluated using the Agilent 2100 High-Sensitivity DNA assay, and concentrations were measured using the Qubit™ dsDNA HS Assay Kit. Libraries were sequenced on a NextSeq 2000 to generate a minimum of 20 million paired-end 60bp reads per sample.

Raw reads were aligned to the mouse reference genome GRCm39 (Ensembl release 106) using the STAR^16^ software v2.7.10a with default parameters. Aligned reads were assigned to genes to give a gene-level count matrix using the featureCounts tool^17^ as part of the Rsubread package v2.10.4. Differential gene expression analysis was performed using the edgeR^18^ software v3.38.1 following the author’s recommended analysis pipeline utilising a quasi-likelihood negative binomial generalized log-linear model.

Gene set enrichment analysis was carried out on fold change ranked gene lists from both male and female mice at 1 week, 3 weeks and 6 weeks post tamoxifen induction of *Il11* expression. The data was imputed into the GSEA software v4.3.2. (Broad Institute, San Diego, USA & UC San Diego, USA) with MSigDB Hallmark gene sets.^19, 20^

KEGG pathway analysis was performed using a combined list of differentially expressed genes across all three-time points. Gene lists were imputed into the ShinyGO v0.77 software (South Dakota State University, USA)^21^ and the top 20 KEGG pathways ranked by enrichment false discovery rate (FDR) values were visualised.

### Protein analysis

Protein extraction was performed using ice-cold Pierce RIPA buffer (ThermoFisher, 89901) supplemented with protease inhibitors (Roche,11697498001) and phosphatase inhibitors (Roche, 4906845001). Tissue was lysed using a Tissue Lyser II (Qiagen) with metallic beads for 3 mins at 30Hz. Protein quantification was performed using a Pierce BCA colorimetric protein assay kit (ThermoFisher, 23225). 10µg of protein was loaded per well and run on a 4- 12% bis-tris precast SDS page gel (Invitrogen, NP0323BOX). Semi-dry transfer was performed using the TransBlot Turbo transfer system (BioRad, 1704150) and the membrane was blocked in 5% BSA (Sigma-Aldrich, A3803). The membrane was incubated overnight at 4°C with primary antibodies phospho-STAT3-Tyr705 (Cell Signalling, 9145S), STAT3 (Cell Signalling, 4904S) and Twist1 (Abcam, ab50887) raised in rabbit. Appropriate secondary HRP-linked antibody was incubated for 1 hour at room temperature and developed using Clarity Western ECL blotting substrate (BioRad, 1705061).

### Masson trichrome staining

Heart and kidney sections were fixed overnight in 10% formalin (Sigma-Aldrich, F5554) and then in ethanol (Fisher Scientific, 10437341, 70% v/v). Samples were embedded in paraffin and staining was performed using the Masson’s Trichrome Kit (Sigma, HT15-1KTKIT) according to the manufacturer’s protocol. Images for analysis were acquired using Zeiss Axio scan digital slide scanner. Quantification was performed by an investigator unbiased of the genotype. Total global collagen was performed using the colour threshold function in the Fiji software to identify fibrotic regions and compared to the whole myocardial area. Perivascular fibrosis was assessed as the degree of fibrosis surrounding vessels as compared to the vessel size. The five largest vessels in each section were analysed and an average was taken for each animal.

### Statistical Analysis

All statistical analyses were performed in GraphPad Prism V9.5.0. Normality testing was performed using the Shapiro-Wilk test. Hypothesis testing for single comparisons was done using an unpaired two ways Student t-test. Comparisons involving male and female mice were performed using a two-way analysis of variance (ANOVA) with Sidak’s multiple comparisons testing. Changes in expression over time were analysed using a one-way ANOVA and changes across genotypes, sex and time in Supplemental Figure 6 were tested using a 3-way ANOVA. All graphs display the mean and standard error of the mean unless stated otherwise. P-values in RNA seq analysis were corrected for multiple testing using the FDR approach. A p-value and FDR of <0.05 was considered significant.

## Results

### IL11 is expressed by cardiomyocytes

TGFβ plays an important role in cardiac pathology following MI and is known to stimulate IL11 secretion from fibroblasts and epithelial cells.^5, 22^ We therefore tested if cardiomyocytes upregulate *Il11* in response to TGFβ stimulation using primary adult mouse cardiomyocytes. As compared to untreated cells, TGFβ stimulation (5ng/mL, 24h) induced a 7-fold increase in the expression of *Il11* (**Fig 1a & b**). To determine the effects of IL11 secretion by cardiomyocytes in the adult *in vivo* we generated a transgenic mouse model (CM-Il11-Tg). The Cre-loxP system using the Rosa26-*Il11* transgenic mouse was used as previously described.^5, 23^ This mouse was crossed with an MCM mouse^24^ **(Fig 1c-e)** to create cardiomyocyte-specific expression of *Il11* in response to tamoxifen administration **(Suppl S1a-c)**.

**Figure 1.**
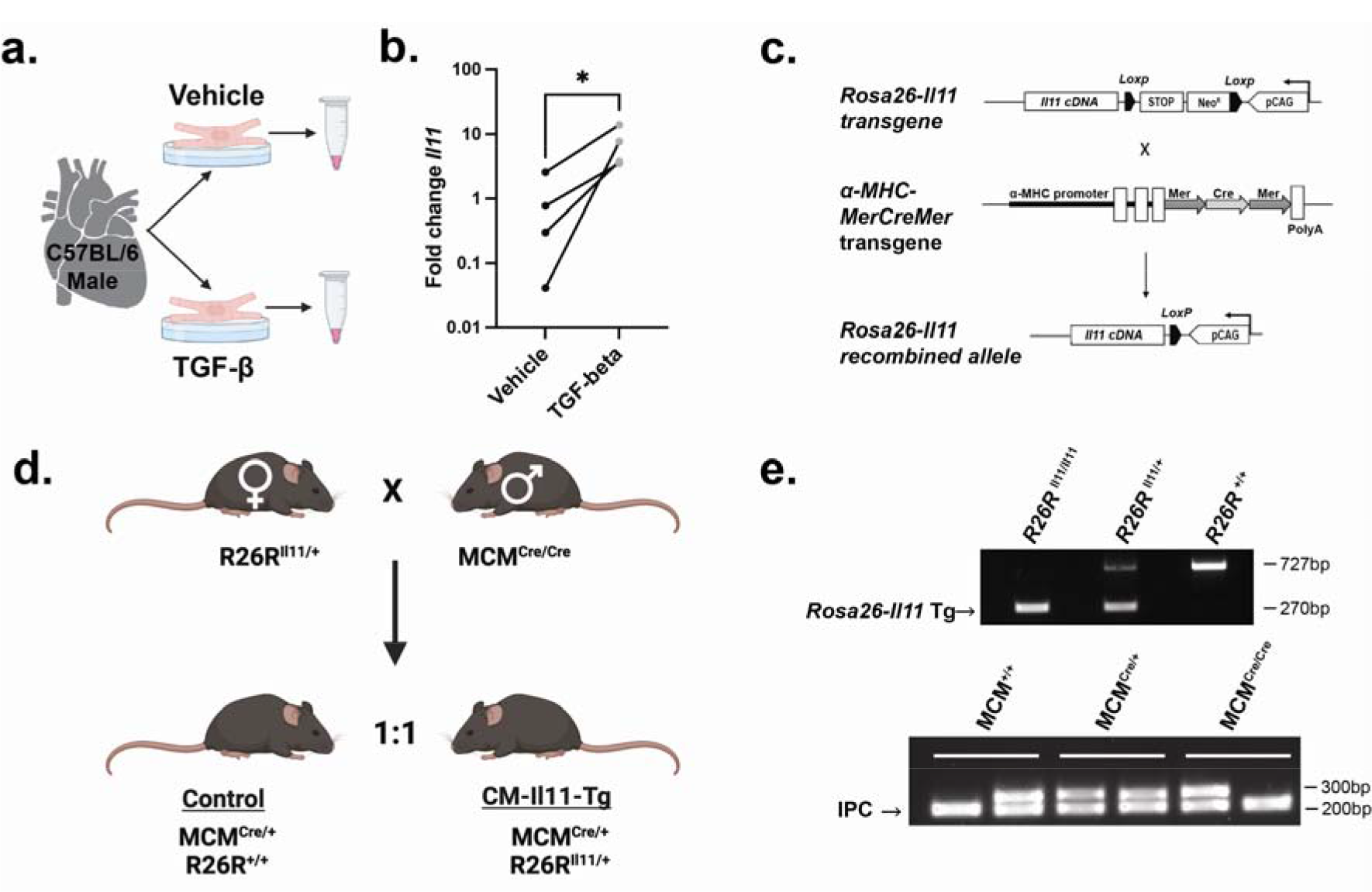
*Il11* expression in cardiomyocytes and generation of a new *Il11* transgenic mouse. (a) Schematic of IL11 of cardiomyocyte extraction and treatment protocol from wild-type C57BL6/J mice. **(b)** QPCR on cell lysates from isolated cardiomyocyte samples after 24 hours of TGFβ treatment (n=4vs4). **(c)** Schematic diagram of the targeted expression of *Il11* in cardiomyocytes. In the *Rosa26-Il11* transgene, a floxed cassette containing both the neomycin (neo) resistance and stop elements is positioned before the murine *Il11* transgene cassette, which undergoes tamoxifen-initiated Cre-mediated recombination when crossed to the α-MHC-modified oestrogen receptor (MCM) mouse (Adapted from Lim et al.).^23^ **(d)** Breeding scheme used to produce CM-Il11-Tg mice and littermate controls. **(e)** Genotyping gels of ear notch biopsy DNA from control and experimental mice. A 270bp band indicated the presence of the Rosa26-*Il11* transgene and the wild-type band appeared at 727 bp (top gel). The MCM band is genotyped using 2 reactions with an internal positive control (IPC). Bands for the wild-type allele or Cre transgene are at 295bp and 300bp respectively. The IPC appears at 200bp. *Stats: Unpaired t-test. P values denoted as *p<0.05*

### IL11 secreted by cardiomyocytes has paracrine profibrotic effects

Following tamoxifen administration, *Il11* was upregulated in the hearts of CM-Il11-Tg mice (**Fig 2a & Suppl S1**). QPCR analysis of myocardial extracts six weeks after tamoxifen administration showed large upregulation of profibrotic (*Col1a1, Col3a1, Fn1, Postn*) and extracellular matrix remodelling (*Mmp2, Mmp9, Mmp14, Timp1*) genes (**Fig 2b & c**). Masson’s trichrome staining of histological sections revealed a large increase in interstitial, epicardial and, most notably, perivascular fibrosis (**Fig 2d-f**). These data suggest paracrine activity of Il11 from cardiomyocytes on fibroblasts to cause cardiac fibrosis that is notably different to the effects of cardiomyocyte-restricted TGFβ expression that does not cause fibrosis of the ventricles.^25^

**Figure 2.**
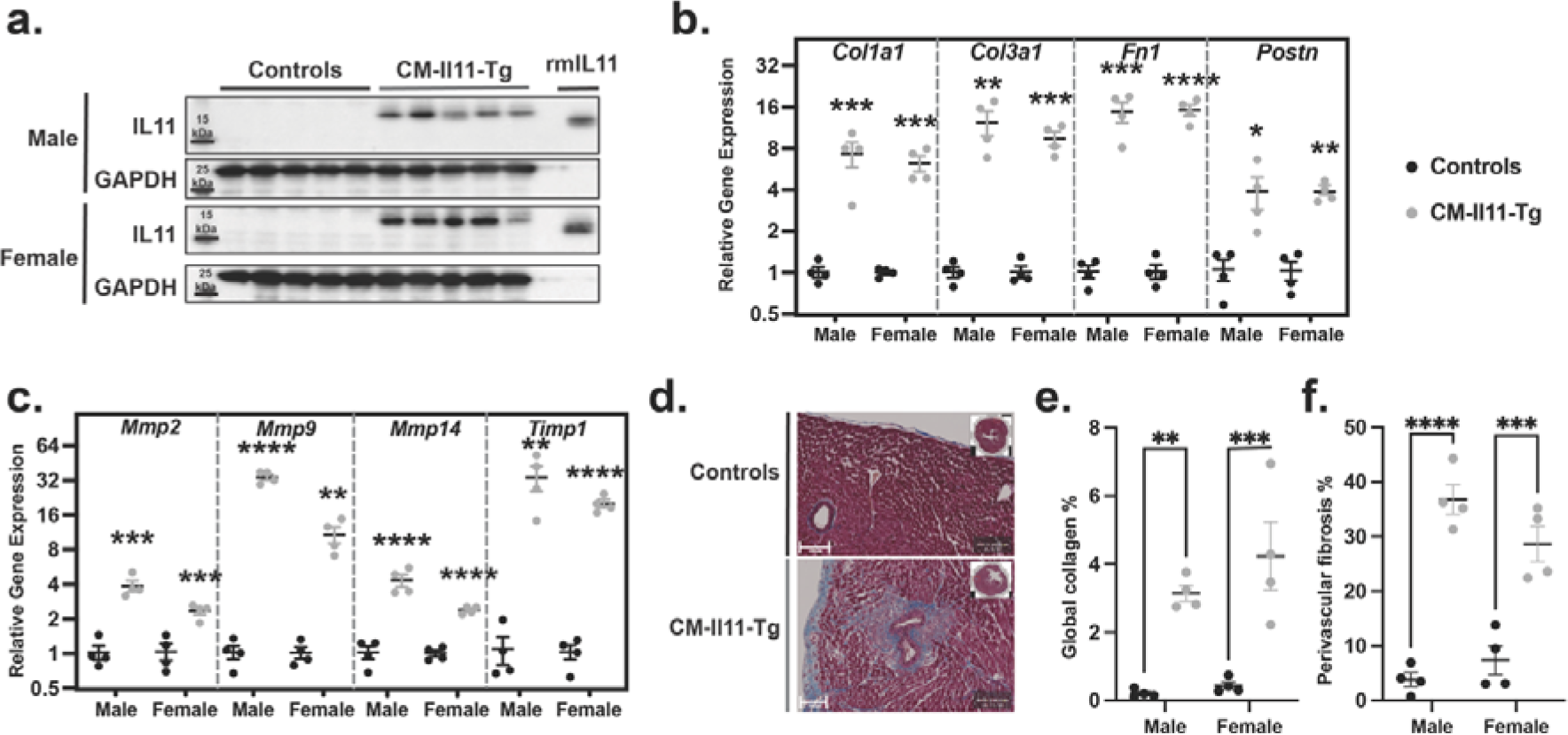
Cardiomyocyte-restricted IL11 expression causes myocardial fibrosis. (a) Ventricular expression of IL11 on western blot 6 weeks after induction of transgene in control and CM-Il11-Tg mice (n=5vs5 mice per sex). QPCR analysis of **(b)** fibrosis and **(c)** extracellular matrix remodelling genes in the ventricular myocardium (n=4vs4). **(d)** Representative Masson’s trichrome staining of control and CM-Il11-Tg mice. Quantification of **(e)** global collagen content of myocardium and **(f)** relative perivascular collagen compared to vessel area. Statistics: *Two-way ANOVA with multiple comparisons*. *p-values donated by *<0.05, **<0.01, ***<0.001,****<0.0001*.

### Proinflammatory effects of IL11 in the heart

RNASeq of left ventricular tissue from male and female mice was performed at 1-, 3- and 6- weeks following induction of IL11 expression and compared to control animals (n=4 per time point per sex). Gene ontology term analysis of CM-Il11-Tg revealed that the top 20 most differentially regulated gene ontology biological processes (GOBP) terms were related to inflammation, immune response, and leukocyte activation (**Fig 3a & b**). These changes were consistent across time points and reproducible in both male and female mice **(Suppl S2&S3)**. Hallmark gene set enrichment analysis (GSEA) also highlighted multiple inflammatory terms in significantly enriched gene sets including Il6 Jak-Stat3 signalling, inflammatory response, interferon-gamma response, TNFα signalling via NFKB and complement activation (**Fig 3d & Suppl S4**).

**Figure 3.**
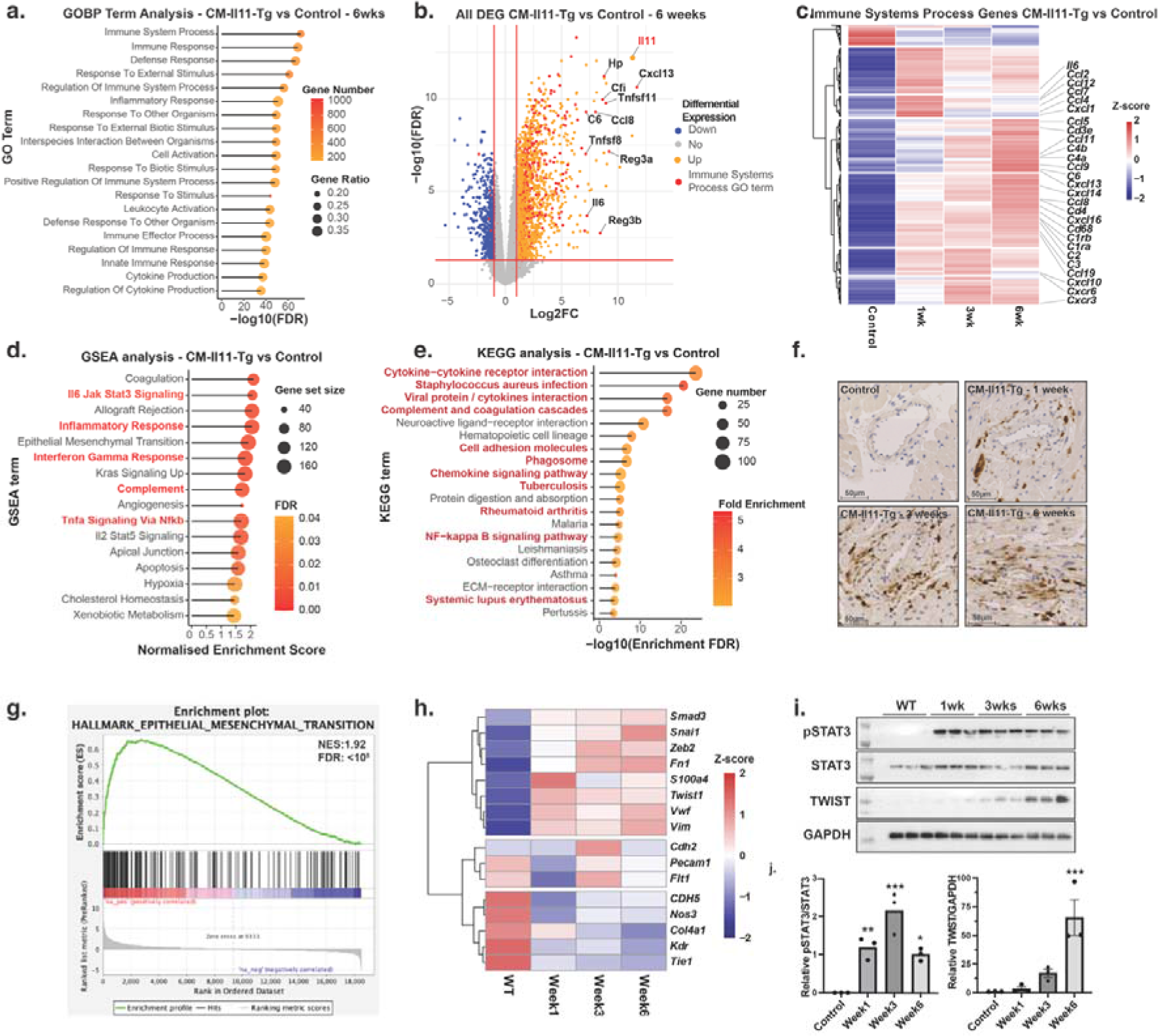
Bulk RNA-seq analysis of the transcriptional effects of IL11 expression in cardiomyocytes. (a) GO term analysis of RNAseq experiment from ventricles of male CM- Il11-Tg mice (n=4) 6 weeks after transgene induction with tamoxifen compared to control animals. **(b)** Volcano plot of all differentially expressed genes (Up=1789 genes, Down=525 genes) in the ventricle from male mice at 6 weeks post transgene induction compared to control animals (n=4vs4). Genes included in the most enriched GOBP term “immune system processes” term are highlighted in red (493/2790 genes) and the top 10 most differentially expressed genes in this GO term are labelled. **(c)** Heatmap with hierarchical clustering of z-scores of differentially expressed “immune system process” genes from ventricular myocardium of male CM-Il11-Tg mice at 1, 3 and 6 weeks after transgene induction compared to control animals. Selected genes involved in leukocyte chemotaxis and the complement system are labelled (n=4 per time point). **(d)** Significantly enriched gene sets from GSEA of 6-week male CM-Il11-Tg mice compared to controls using MSigDB Hallmark gene sets (n=4vs4). **(e)** Top 20 most significantly enriched KEGG terms on pooled analysis of all differentially expressed genes across 3 time points in male mice (n=4 per time point). KEGG terms involved in the inflammatory response are highlighted in red. **(f)** Immunohistochemical staining of macrophages using Mac2 antibody in the perivascular region of control mice and CM-Il11-Tg male mice 1, 3 and 6 weeks after transgene induction. **(g)** Enrichment plot of epithelial to mesenchymal transition (EMT) gene set (M5930) using Broad GSEA software 6 weeks after induction of the transgene in CM-Il11-Tg mice compared to control mice (n=4) (NES:1.92, FDR: <0.0001) (h) Heatmap of z-scores of genes involved in endothelial to mesenchymal transition including mesenchymal markers, endothelial markers and related transcription factors at 1, 3 and 6 weeks post-tamoxifen induction. **(i)** Western blot and quantification of phosphorylated STAT3, total STAT3 and TWIST1 expression in the ventricular myocardium in control mice and CM-Il11-Tg mice 1, 3 and 6 weeks after induction of *Il11* expression (n=3 per time point).

Hierarchical clustering of differentially expressed genes from the most enriched GOBP term “immune system process” (GO:0002376) demonstrates clusters of gene expression vary across the individual time points. In particular, genes associated with major histocompatibility complexes are significantly upregulated at 3 and 6 weeks compared to week 1 (*H2-Ab1, H2-DMb1, H2-Eb1*) which is suggestive of infiltration and activation of phagocytic antigen-presenting cells into the myocardium. Additional genes which upregulated at later time points include factors within the complement pathway (*C4a, C4b, C6),* immunoglobulin-related genes (*Igf2, Ighg2c* and *Jchain)* and lymphocyte markers and chemotactic signalling molecules (*CD4, Cd3e* and *Cxcl13)*. A smaller cluster of genes most strongly upregulated at the one-week time point included chemokines involved in the recruitment and activation of granulocytes and monocytes (*Il6, Tnf, Cxcl1, Cxcl5, Ccl4, Ccl7*) **(Fig 3c)**.

KEGG pathway analysis is consistent with the GOBP analysis and includes complement activation, phagosome activity, chemokine signalling pathways and the NF-kappa B signalling pathway among the most significantly enriched pathways **(Fig 4e & Suppl S5)**. Immunohistochemical staining for macrophage supports the development of a proinflammatory environment within the myocardium of CM-Il11-Tg mice with a focus of inflammatory changes in the perivascular area (**Fig 3f).**

**Figure 4.**
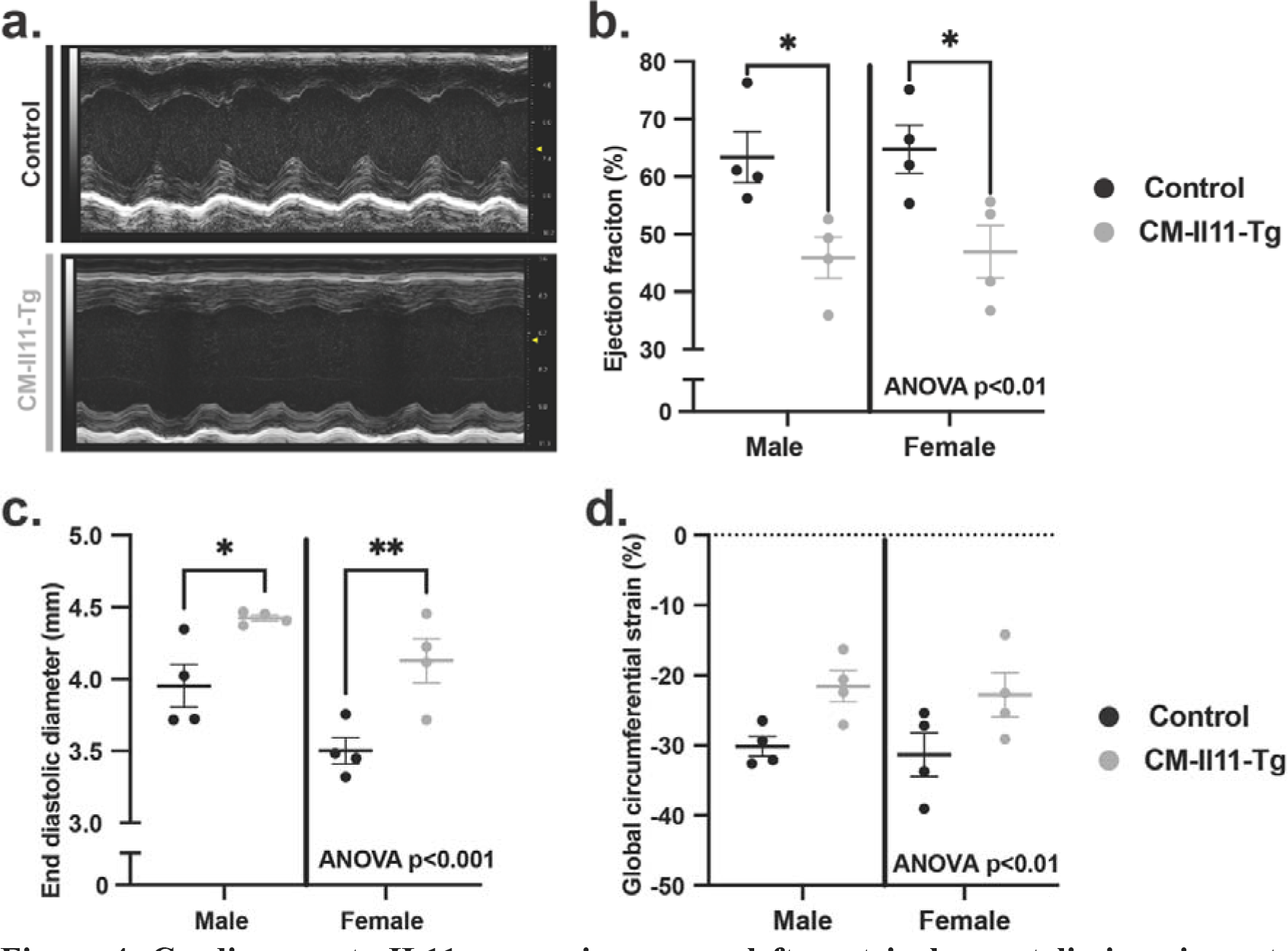
Cardiomyocyte IL11 expression causes left ventricular systolic impairment. (a) Representative M-mode images from parasternal long axis (PLAX) male control and CM- Il11-Tg mice 6 weeks after induction of *Il11* expression. **(b)** Quantification of left ventricular ejection fraction from PLAX M-mode images in male and female CM-Il11-Tg mice compared to control animals 6 weeks after transgene induction (n=4vs4 per sex). **(c)** End diastolic diameter measured from PLAX M-mode images (n=4vs4 per sex). **(d)** Global circumferential strain measured using Vevostrain speckle tracking software from the parasternal short axis view (n=4vs4 per sex). *Stats: Two-way ANOVA with multiple comparisons. p-values denoted by *<0.05, **<0.01*.

### Endothelial-to-mesenchymal transition in IL11 expressing mice

Beyond fibrosis and inflammation, IL11 has been implicated in epithelial-to-mesenchymal transition (EMT) and endothelial-to-mesenchymal transition (EndMT), which may be of relevance to the strong perivascular phenotype we observed following *Il11* expression (**Fig 2d**).^6, 26^ The EMT Hallmark geneset was in the top six most enriched genesets following transgene recombination across all time points in male and female mice **(Fig 3d, g & Suppl S4).** Reciprocal to upregulation of EMT genes and mesenchymal genes in CM-Il11-Tg mice, endothelial markers (*Cdh5, Kdr, Flt1 and Tie1 and Nos3*) were downregulated **(Fig 3h).** Twist a key transcription factor in EndMT and EMT was upregulated at the protein level at later time points in the CM-Il11-Tg mice **(Fig 3i).**

### Impaired ventricular function following cardiomyocyte IL11 expression

Transthoracic echocardiography performed on CM-Il11-Tg mice 6 weeks after transgene induction showed similar impairment of systolic function in both male and female CM-Il11- Tg mice, as compared to tamoxifen injected controls **(Fig 4a & b)** suggesting the absence of sexual dimorphism of cardiomyocyte-specific *Il11* expression on heart function. This was accompanied by dilation of the left ventricular cavity and a reduction in global circumferential strain in CM-Il11-Tg mice. Systolic dysfunction was observed within 3 weeks of transgene induction with differences in diastolic LV size apparent only at the later 6-week time point (**Suppl S6**).

## Discussion

Until recently IL11 was thought to be anti-fibrotic in the heart and other organs and is still thought by many as anti-inflammatory, although these accepted concepts have been challenged by studies in recent years.^2, 5, 6, 8, 27–29^ Here we examined the specific effect of IL11 secreted from cardiomyocytes, which we show to express *Il11* in response to TGFβ stimulation. While TGFβ is viewed as the strongest profibrotic factor, it was striking that cardiomyocyte-restricted *Il11* expression caused robust ventricular fibrosis whereas similar expression of TGFβ in the same experimental system does not.^25^ This difference may reflect the proinflammatory effects of IL11 in a maladaptive loop of fibro-inflammation, whereas TGFβ is anti-inflammatory.^30–32^ The data may also reflect that the paracrine effects of cardiomyocyte-derived lL11 on neighbouring fibroblasts are more pronounced than those of TGFβ, which might be more important as an autocrine factor in fibroblasts.

In our studies, the cardiac inflammation seen following *Il11* induction in mouse cardiomyocytes evolved over time with an initial cytokine signature and a later footprint of chemokines and complement factors. These effects were profound and will reflect, in part, the recruitment of neutrophils, leukocytes and monocytes, which were apparent on Mac2 staining. While a review in 2022^2^ speculated that the jury was still out as to whether IL11 was pro- or anti-inflammatory, we think these data, taken together with other recent publications, strongly support the idea that IL11 is proinflammatory.^2, 29^ Our data also show that IL11 causes cardiac dysfunction, as seen with chronic recombinant IL11 injection to the mouse and IL11 expression in the fibroblast compartment, which further consolidates the negative impact of IL11 on cardiac function.^5^

We noted that IL11 expression induced extensive changes within the perivascular region of the myocardium and that this was associated with reduced expression of endothelial markers and upregulation of EndMT genes. Intriguingly, IL11 was found to have a role in EndMT in the pulmonary arteries in the setting of idiopathic pulmonary fibrosis-related pulmonary hypertension.^6^ We propose that IL11-driven EndMT, directly or indirectly, contributes to the perivascular fibrosis demonstrated here and in pulmonary hypertension. Our findings of IL11-associated EMT/EndMT are in keeping with recent studies in kidney disease where IL11 causes pathogenic EMT.^4^

This study has limitations. It is likely that tissue levels of IL11 produced following transgene recombination are higher than those seen in pathology. However, this expression system provides valuable insights into the cardiac biology associated with gene gain-of-function and has been used extensively, including previously to study TGFβ effects on cardiac fibrosis, which were minimal. Additionally, we have not been able to ascertain to what degree inflammation in the heart is due to infiltration of immune cells or activation of resident cells. Irrespective of the origin, increased inflammation driven by either mechanism would be expected to adversely impact cardiac function. The endMT changes that we observed do not exclude the possibility that endothelial cell signatures are reduced due to endothelial cell loss from another mechanism; the concurrent expression of pro-EMT transcription factors strengthens our interpretation.

We suggest that cardiomyocyte stress that occurs with ischaemia or pressure overload^10, 11^ results in the secretion of IL11 from cardiomyocytes which leads to cardiac fibrosis and inflammation. Cardiac fibrosis remains an attractive therapeutic target but unfortunately, inhibition of TGFβ failed in the clinic due to on-target toxicities, which include inflammation and cardiac impairment.^33, 34^ IL11 may therefore represent an alternative target as its effects on cardiac fibrosis appear more profound than TGFβ and IL11 is also proinflammatory. Mice deleted for *Il11* or *Il11ra1* or administered anti-IL11 antibody, appear to have no major health issues indicating a favourable safety profile, again contrasting with genetic or pharmacologic inhibition of TGFβ that is detrimental.^30, 33^ To conclude, our data show IL11 to be profibrotic and proinflammatory in the heart and we propose IL11 as a potential therapeutic target for fibro-inflammatory heart disease.

## Acknowledgements

For the purpose of open access, the authors have applied a Creative Commons Attribution (CC BY) licence to any Author Accepted Manuscript version arising. Figures were created with BioRender.com

## Abbreviations

ANOVA: Analysis of variance
CM-Il11-Tg: Cardiomyocyte specific Il11 expression transgenic
EMT: Epithelial to mesenchymal transition
EndMT: Endothelial to mesenchymal transition
FDR: False discovery rate
GOBP: Gene ontology biological process
GSEA: Gene set enrichment analysis
IL11: Interleukin 11
MHC: Myosin heavy chain
MerCreMer: Modified estrogen receptor
PLAX: Parasternal long axis
rhIL11: Recombinant human interleukin 11
rmIL11: Recombinant mouse interleukin 11
TGFβ: Transforming growth factor β

