## Supplemental Materials for "Cardiomyocyte-restricted expression of IL11 causes cardiac fibrosis, inflammation, and dysfunction"

### **Supplementary Figures**

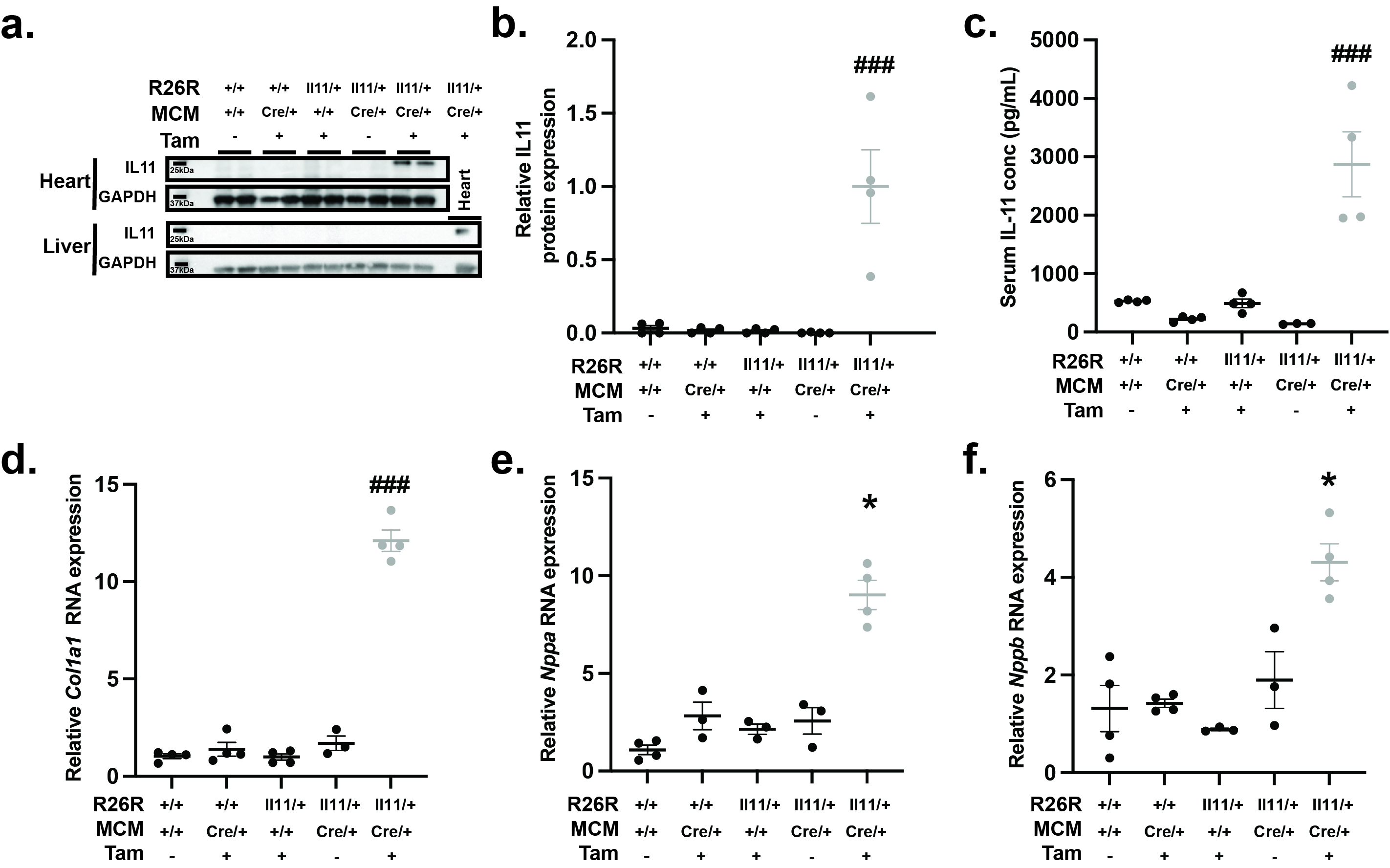

**Supplementary S1. (a)** Western blot of IL11 expression in the ventricular myocardium and liver in male mice with different combinations of R26R-Il11 transgene, MCM gene and tamoxifen administration. Demonstrating myocardial-specific expression in response to tamoxifen administration without detectable hepatic expression. (b) Quantification of western blot from (a) (n=4 per condition) (c) Serum IL11 concentration. (d-f) QPCR of myocardial tissue from male mice targeting *Col1a1, Nppa and Nppb* in combinations of genotypes and tamoxifen administration (n=4 per condition).  *P-values denoted by: *p<0.05 vs R26R^+/+^:MCM^+/+^ in the absence of tamoxifen. ### p<0.001 vs all other groups. Stats: One-way ANOVA with multiple comparisons.*

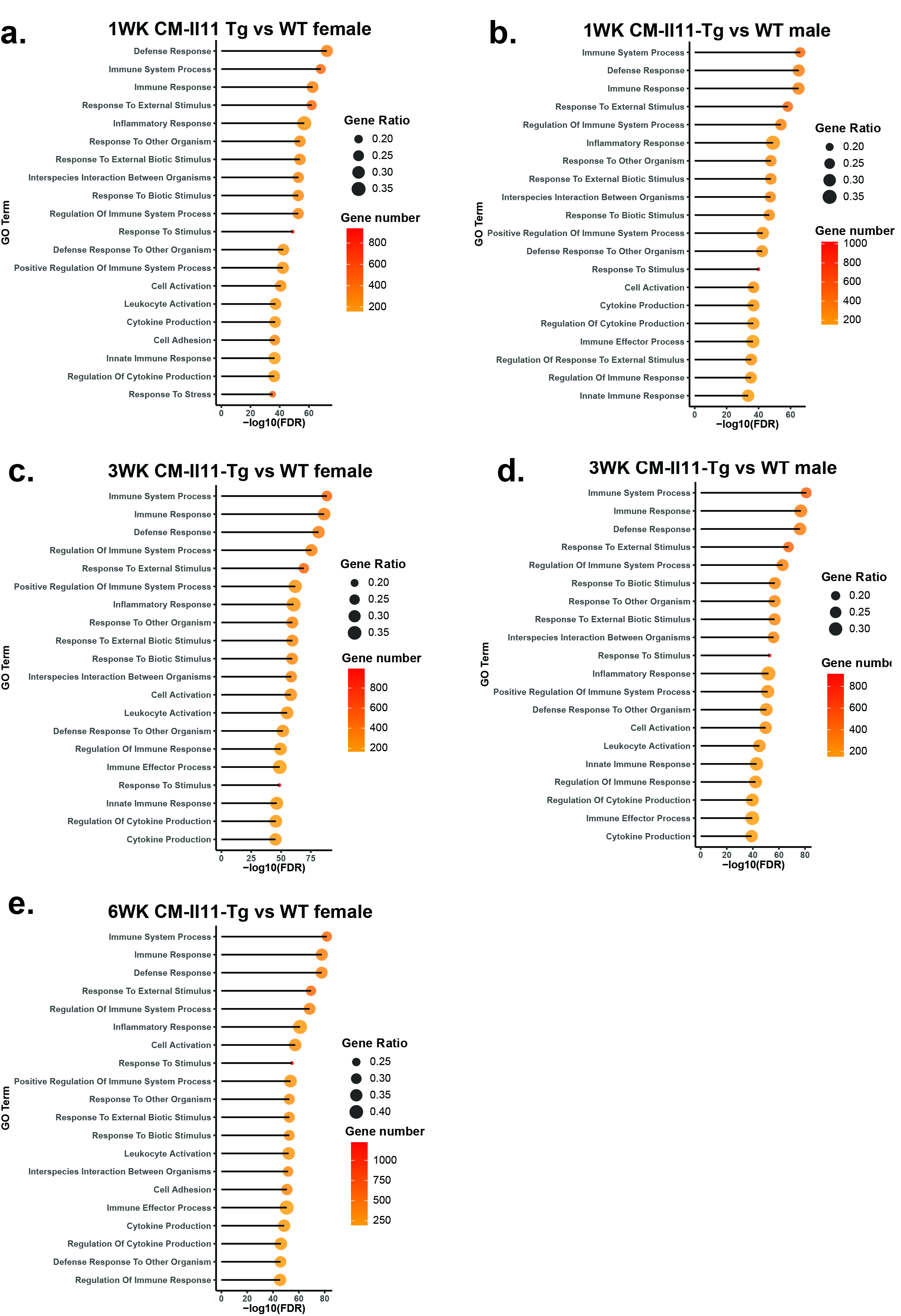

**Supplementary S2. Top 20 Biological processes GO terms.** Top 20 gene ontology biological processes terms from gene ontology analysis of differentially expressed genes in male and female CM-Il11-Tg mice compared to controls at 1, 3 and 6 weeks after administration of tamoxifen.

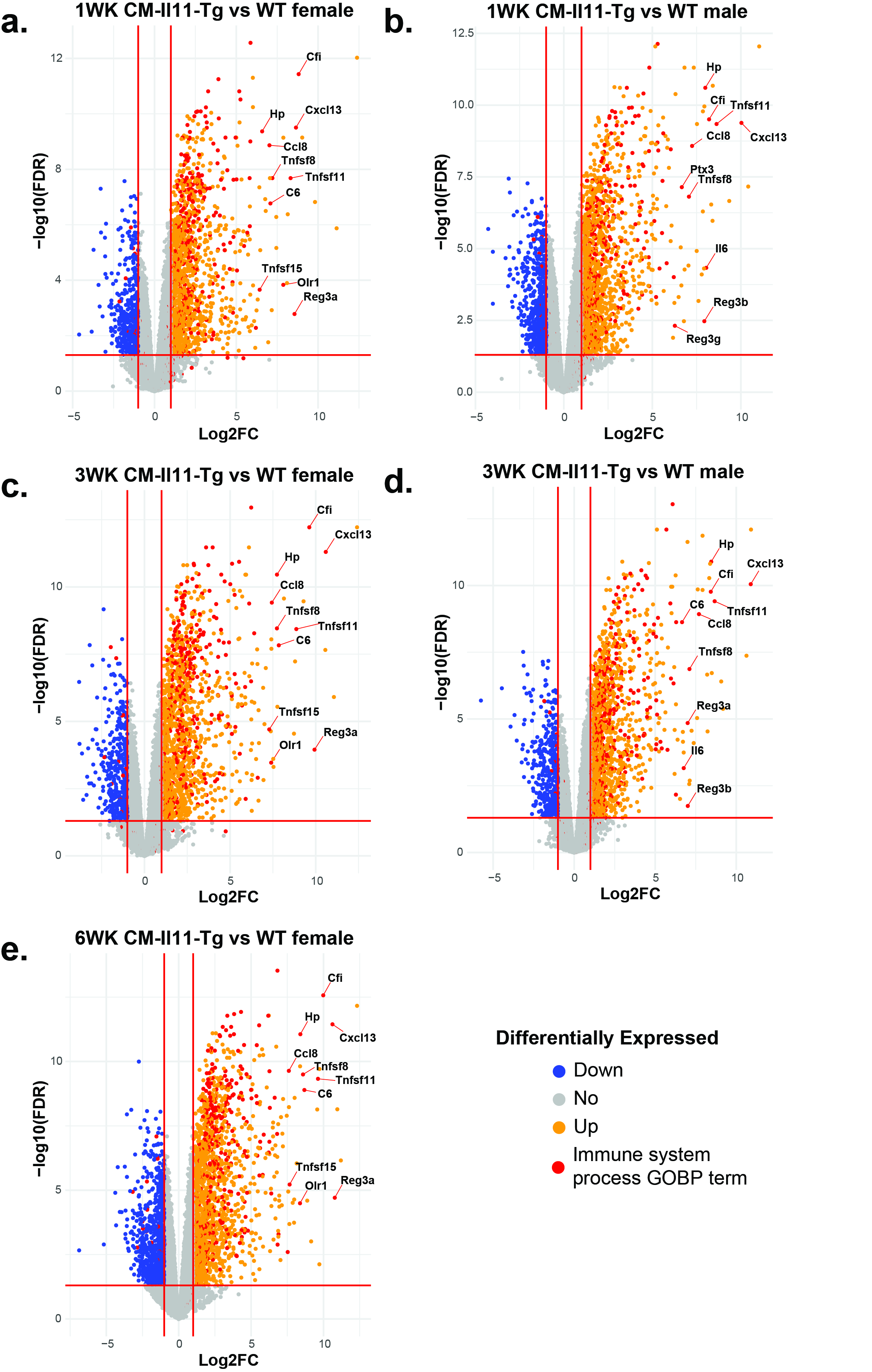

**Supplementary S3. Volcano plot of differentially expressed genes.** Visualisation of all detected genes in myocardial RNA seq data in males and female CM-Il11-Tg mice compared to controls at 1, 3 and 6 weeks after administration of tamoxifen. Red lines are placed at log2Fc of 1 or -1 and at a false discovery rate of 0.05. Genes involved in the most enriched GOBP term “immune system processes” are highlighted in red and the top 10 genes in this GO term are labelled (n=4vs4 per time point).

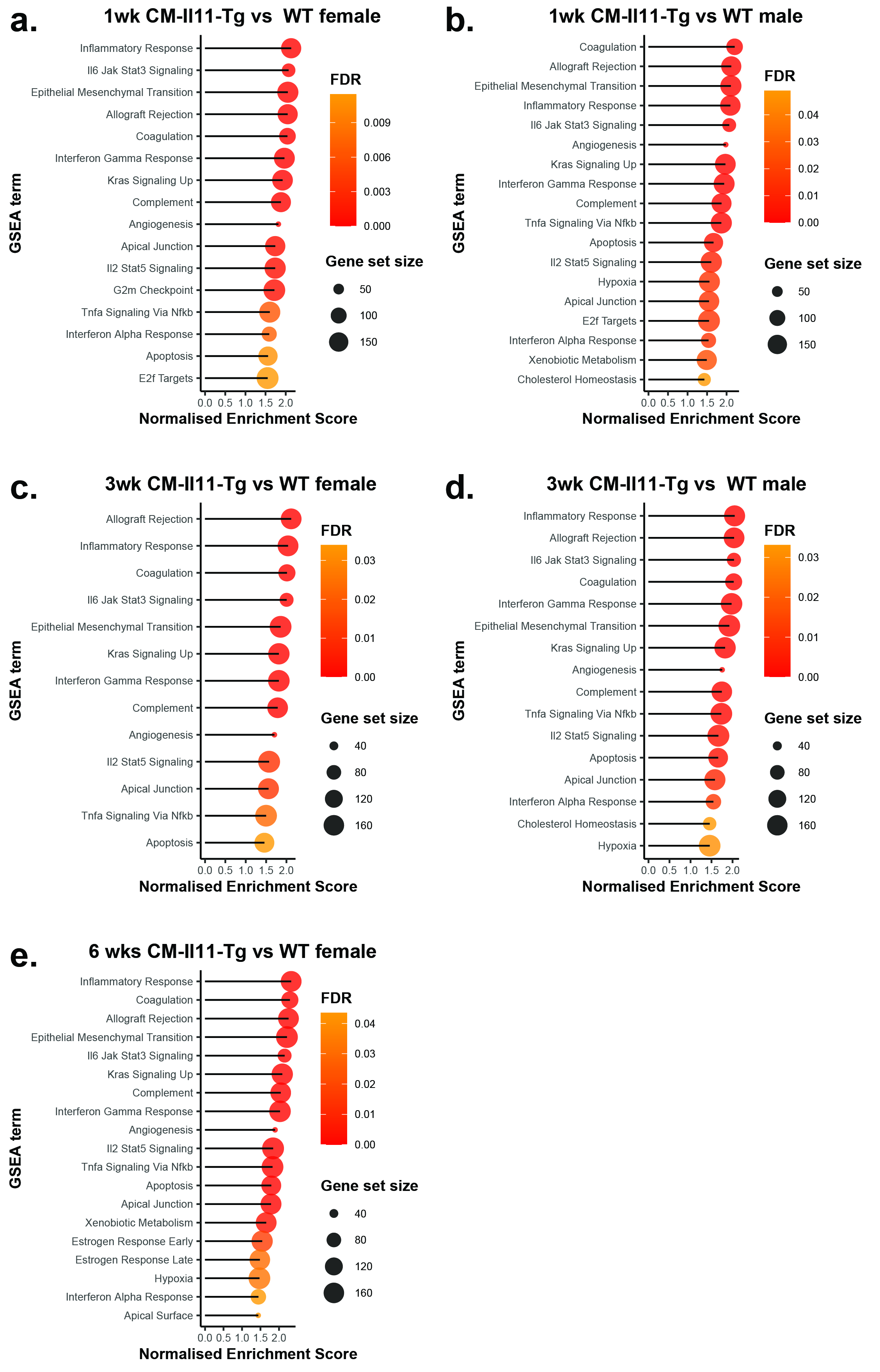

**Supplementary S4. Gene set enrichment analysis using Hallmark gene sets.** Upregulated hallmark gene sets using GSEA in male and female CM-Il11-Tg mice compared to controls at 1, 3 and 6 weeks after tamoxifen administration.

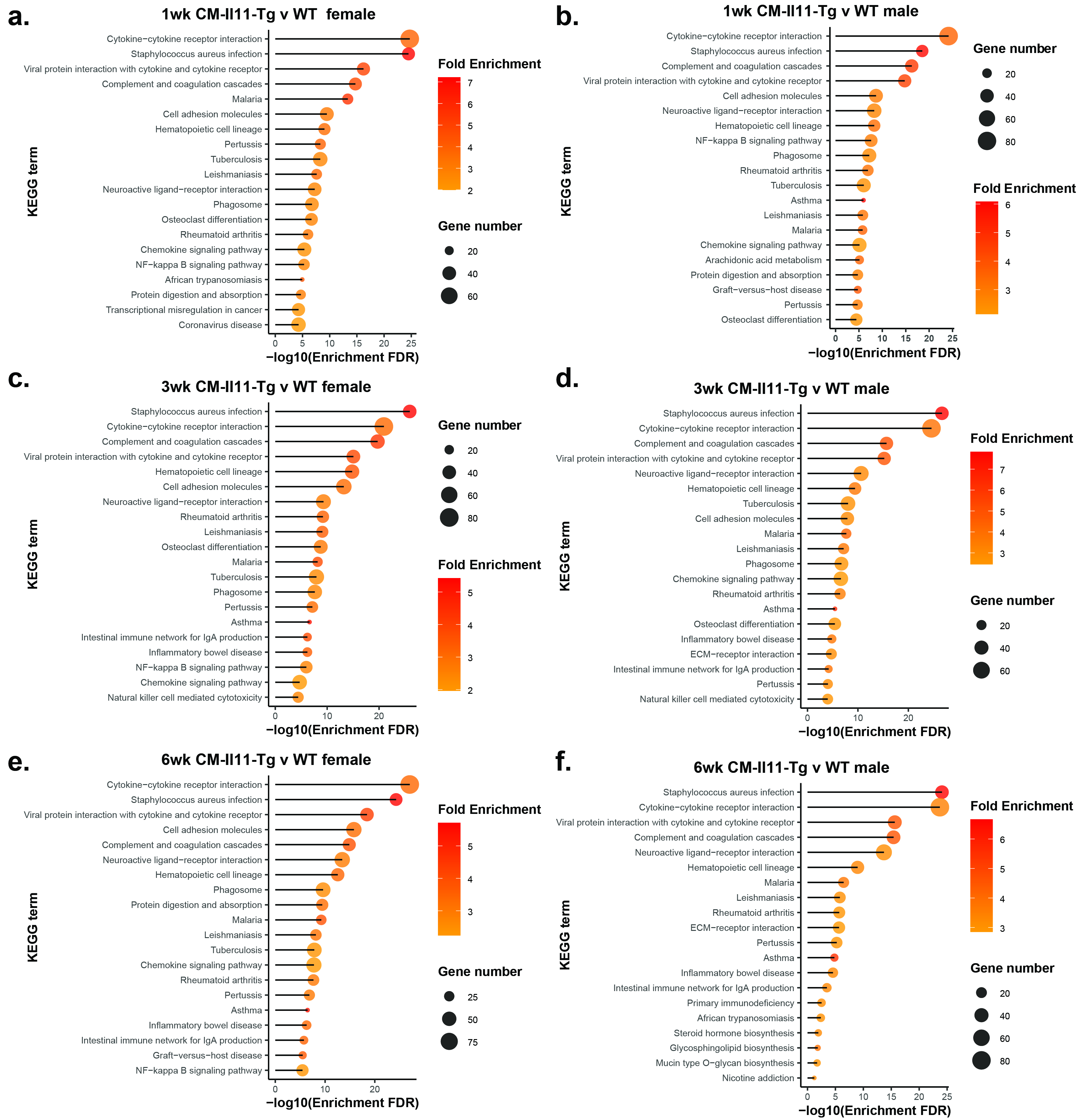

**Supplementary S5. KEGG analysis at individual time points.** Top 20 most significantly enriched KEGG terms based on differentially expressed genes in male and female CM-Il11-Tg mice compared to controls at 1, 3 and 6 weeks after tamoxifen administration.

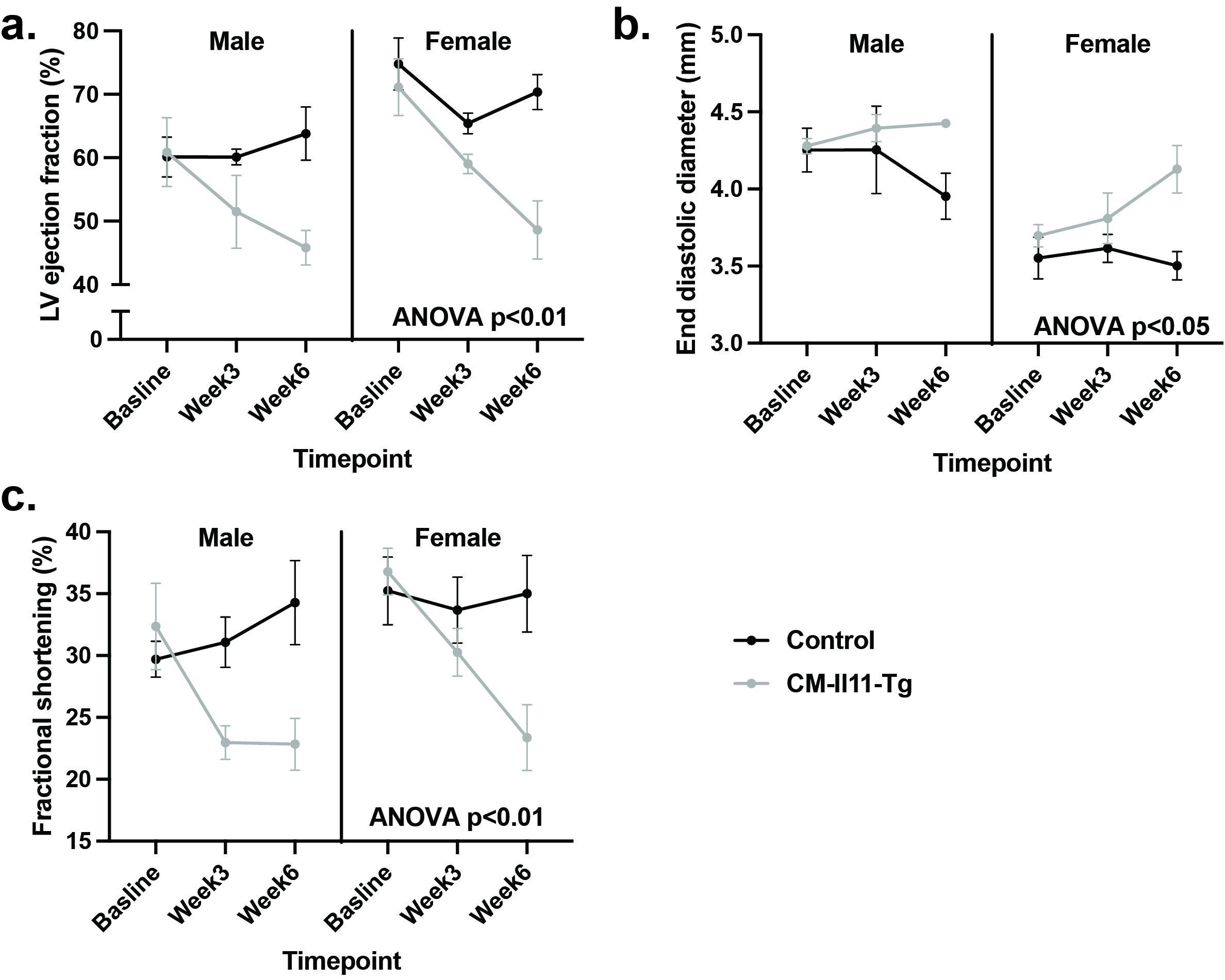

**Supplementary S6. Cardiac function changes compared to baseline at 3 weeks and 6 weeks. (a)** Left ventricular ejection fraction in male and female CM-Il11-Tg mice at baseline, 3 and 6 weeks after transgene recombination. **(b)** End diastolic volume at baseline, 3 and 6 weeks. **(c)** Fractional shortening at baseline, 3 and 6 weeks. (n=4 per group per sex) *Stats: 3 way ANOVA*

### **Supplementary Tables**

| **Rosa26*-Il11*** | **Forward** | **GTTTTGGAGGCAGGAAGCACTTGC** |
| --- | --- | --- |
|  | **Forward** | **GCAGTGAGAAGAGTACCACCATGAGTCC** |
|  | **Reverse** | **CAATGCTCTGTCTAGGGGTTGGATAAGC** |
| **α-MHC-MerCreMer** | **Forward** | **5'-TCTATTGCACACAGCAATCCA-3'** |
|  | **WT reverse** | **5'-CCAACTCTTGTGAGAGGAGCA-3'** |
|  | **Mutant reverse** | **5'-CCAGCATTGTGAGAACAAGG-3'** |

**Supplementary table 1. Primers used for genotyping PCR.**

| **Target** | **Taqman primer/probe** |
| --- | --- |
| *Col1a1* | Mm00801666_g1 |
| *Col3a1* | Mm00802300_m1 |
| *Fn1* | Mm01256744_m1 |
| *Il11* | Mm00434162_m1 |
| *Mmp2* | Mm00439498_m1 |
| *Mmp9* | Mm00442991_m1 |
| *Mmp14* | Mm00485054_m1 |
| *Postn* | Mm01284919_m1 |
| *Timp1* | Mm01341361_m1 |
| *Gapdh* | Mm99999915_g1 |

**Supplementary table 2.** Taqman probes used for qPCR
